# Reliably Measuring Learning-Dependent Distractor Suppression with Eye Tracking

**DOI:** 10.1101/2024.02.23.581757

**Authors:** Andy J. Kim, Laurent Grégoire, Brian A. Anderson

## Abstract

In the field of psychological science, behavioral performance in computer-based cognitive tasks often exhibits poor reliability. The absence of reliable measures of cognitive processes contributes to non-reproducibility in the field and impedes investigation of individual differences. Specifically in visual search paradigms, response time-based measures have shown poor test-retest reliability and internal consistency across attention capture and distractor suppression, but one study has demonstrated the potential for oculomotor measures to exhibit superior reliability. Therefore, in this study, we investigated three datasets to compare the reliability of learning-dependent distractor suppression measured via distractor fixations (oculomotor capture) and latency to fixate the target (fixation times). Our findings reveal superior split-half reliability of oculomotor capture compared to that of fixation times regardless of the critical distractor comparison, with the reliability of oculomotor capture in most cases falling within the range that is acceptable for the investigation of individual differences. We additionally find that older adults have superior oculomotor reliability compared with young adults, potentially addressing a significant limitation in the aging literature of high variability in response time measures due to slower responses. Our findings highlight the utility of measuring eye movements in the pursuit of reliable indicators of distractor processing and the need to further test and develop additional measures in other sensory domains to maximize statistical power, reliability, and reproducibility.

## Introduction

The field of psychological science was challenged in the past decade to improve the replicability of behavioral research based on large scale examples of non-reproducibility (Johnson et al., 2017; Open Science Collaboration, 2012, 2015). Nosek and colleagues define reproducibility, robustness, and replicability as “testing the reliability of a prior finding” and propose that maximizing the reliability of research findings will improve research credibility and the translation of knowledge into application (Nosek et al., 2022). The reliability of measurements is particularly important when maximizing the power of significance tests, and measures with poor reliability are not sensitive in detecting individual differences (Zimmerman et al., 1993). Researchers have commonly utilized two types of measurements of reliability: test-retest reliability and internal consistency (split-half correlation). These tests have often revealed poor reliability of behavioral measures in the field of psychological science (Dang et al., 2020; Draheim et al., 2019; Paap & Sawi, 2016), calling researchers in the field to identify and develop more reliable measures that can be consistent across multiple experimental paradigms.

As the critical need for reliable measures is increasingly recognized, researchers utilizing visual search paradigms have recently highlighted the poor reliability of measures specifically using behavioral response times. Ivanov et al. (2023) investigated whether difference scores in manual response times and accuracy were reliable and could be utilized as an individual-level measure. Utilizing both split-half and test-retest reliability measurements, the authors investigated whether attention capture, learned distractor suppression at a high-probability location in the visual search array, and corresponding suppression of targets at the high-probability location could serve as reliable measures for investigating individual differences (Ivanov et al., 2023). Over the three measures, the authors report poor to moderate split-half reliability over response times and poor reliability over accuracy, in addition to poor test-retest reliability with respect to both response times and accuracy. Furthermore, three studies investigating selection history effects of reward learning in visual search also reported poor test-retest reliability of behavioral response times (Anderson & Kim, 2019; Freichel et al., 2023; Garre-Frutos et al., 2024). These studies collectively identified that response time exhibits poor reliability over experience-driven attention effects. However, in Anderson and Kim (2019), value-driven oculomotor capture exhibited strong test-retest reliability, suggesting that oculomotor capture may be more sensitive and reliable in contrast to oculomotor fixation times and even more so when compared with manual response times (Anderson & Kim, 2019; Weichselbaum et al., 2018).

Therefore, in the current study, we investigated whether oculomotor measures of distractor fixations provide superior reliability compared to response time-based measures (fixation time or time to make an eye movement to the target). We investigated oculomotor measures in three studies containing a total of 8 experiments that utilized a visual search task incorporating attention capture and/or distractor suppression. The selected studies were limited to investigating the reliability of distractor suppression in the context of selection history effects, given pessimistic findings concerning manual response time measures (Ivanov et al., 2023). We aimed to examine the reliability of oculomotor measures in visual search across multiple experimental paradigms incorporating statistical learning of a high-probability distractor location, learned value-associations with the distractor in a context in which these associations lead to reduced distractor interference, and proactive distractor suppression (feature-search) vs. reactive distractor disengagement (singleton-search). Thus, we look to evaluate the reliability of oculomotor measures across numerous critical distractor comparisons. In two cases, data from both older and younger adults was available, permitting an assessment of the reliability of oculomotor measures as a function of age. Based on the findings of Anderson and Kim (2019), we hypothesize that the reliability of oculomotor capture measures will be superior to that of measures involving fixation time, and that these oculomotor measures will also demonstrate high reliability that is superior to the characteristically low reliability associated with manual response time measures as observed in the literature.

## Methods

### Datasets

We evaluated three datasets that incorporated oculomotor measures in visual search tasks to investigate the reliability of oculomotor capture by the distractor and fixation times (oculomotor response times) between two critical distractor conditions (Grégoire et al., 2022; Kim et al., 2024; Kim & Anderson, 2022). In Kim and Anderson (2022), the critical distractor comparison was a distractor appearing at a high-probability location vs. a distractor appearing at a low-probability location (statistical learning of a high-probability distractor location). In Grégoire et al. (2022), the critical distractor comparison was previously conditioned distractors (CS+; associated with reward or electric shock) vs. neutral distractors (value- and threat-modulated attentional capture). In this latter study, we separated findings over the three experiments (focusing on the first two in which distractor suppression was observed). In Kim et al. (2024), the critical distractor comparison was attention capture by the distractor on distractor-present trials (first saccade to the distractor) vs. first fixation to a single non-target in distractor-absent trials (attention capture by a physically salient distractor when engaging in feature-search or msingleton-search mode); reliability scores were separated by both experiments (feature-search vs. singleton-search) and calculated separately among young and older adult samples to probe potential age differences.

### Split-Half Reliability

Instead of utilizing an arbitrary odd vs. even split, we estimated internal consistency by utilizing a permuted random split procedure as in Garre-Frutos et al. (2024). In this procedure, all trials were randomly split into two halves with an equal number of observations in each half per condition per run to account for time-dependent effects (e.g., learning or extinction). Trials for each half were then concatenated over all runs. Then, a difference score between the two critical distractor conditions was computed for each concatenated half for each participant and correlated to get a Pearson’s *r* correlation coefficient. This procedure was repeated 1000 times and the correlation coefficients were averaged to compute the mean split-half correlation.

### Non-Parametric Randomization Tests

To determine whether estimates of reliability for oculomotor capture and fixation times were significantly different across conditions, we conducted non-parametric randomization tests. Based on the 1000 split-half correlation coefficients calculated for each measure (before averaging), we first computed the mean of the difference scores between the oculomotor capture and fixation time measures as the true sample mean. Then, from the combined 2000 coefficient values for both measures, we randomly assigned 1000 values to each measure to create two unique sample groups and computed the difference of these group mean *r* values (random sample), under the null hypothesis that there was no difference between split-half reliability obtained using each measure and, thus, random assignment of reliability to a dependent measure should tend to produce a similar difference score to the difference score observed between the two measures in the actual data. This randomization procedure was repeated 1000 times and the p-value was manually calculated from the z-score using the observed sample mean.

## Data Availability

The datasets analyzed in the current study are publicly available in the Open Science Framework repository, https://osf.io/fkj92/.

## Results

### Kim and Anderson (2022)

In Kim and Anderson (2022), visual search required fixating on a target shape singleton in the absence and presence of a salient color singleton distractor. Critically, the location of the color distractor in distractor-present trials was in a high-probability location 45% of the time and equally often in the other low-probability locations (5 low-probability locations). When comparing the oculomotor measures, the split-half correlation for the learning-dependent reduction in oculomotor capture (probability of fixating the distractor on low-probability minus high-probability trials) was *r* = .802 and for fixation time (latency to fixate the target on low-probability minus high-probability trials) was *r* = .698. Using non-parametric randomization tests, we found that the reliability of oculomotor capture was significantly superior compared to fixation time, *p <* .001 (see Figure 1).

**Figure 1.**
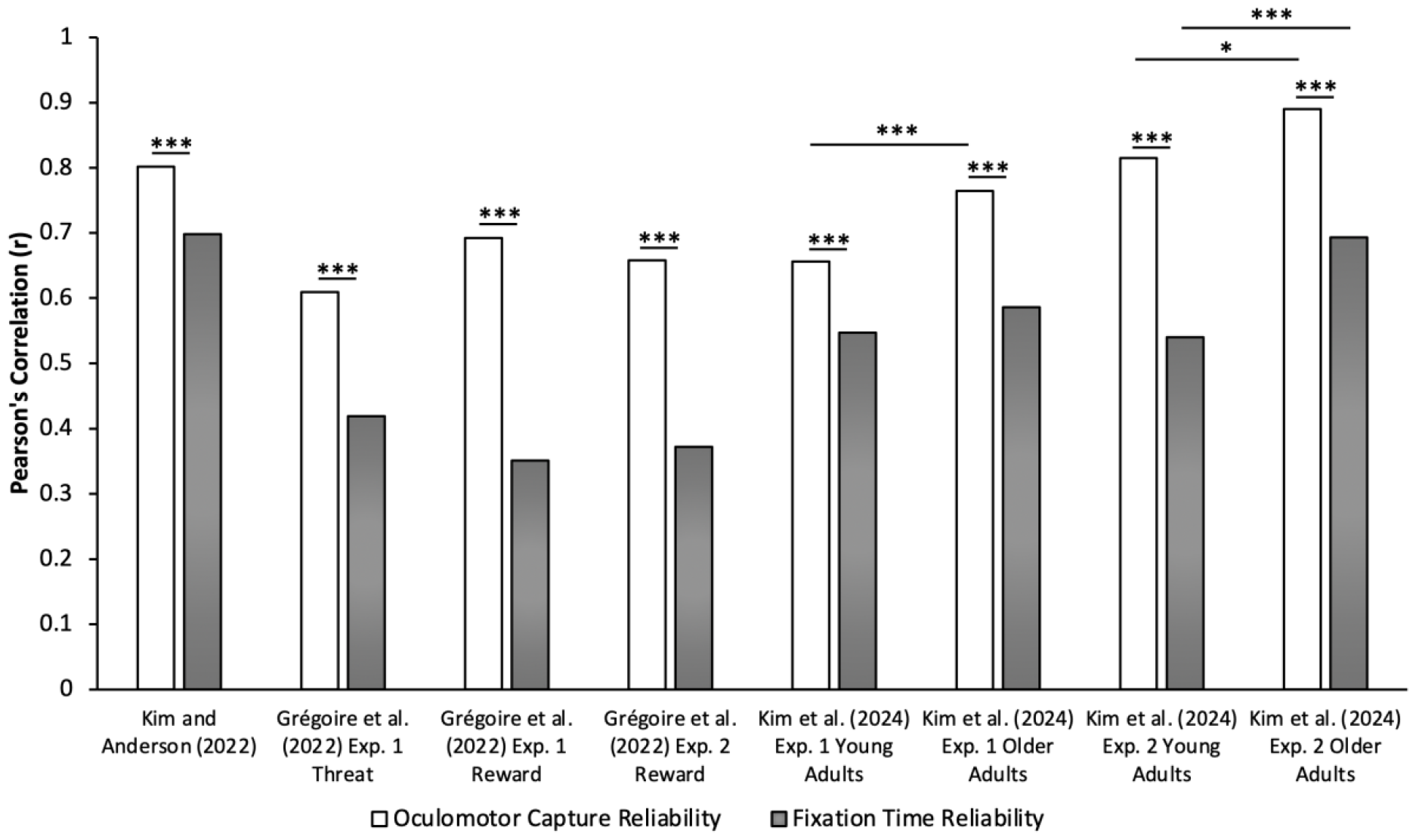
Split-half reliability of oculomotor capture is superior to reliability of fixation times. Bar graphs depict Pearson’s correlation values over attention capture by the distractor (oculomotor capture) and fixation times across multiple datasets. Regardless of critical distractor comparisons (high-vs. low-probability location; reward/threat-related vs. neutral; distractor-present vs. distractor-absent), type of visual search attentional template (feature-search vs. singleton-search), and age groups (young adults vs. older adults), the reliability of oculomotor capture was superior to the reliability of fixation times. Furthermore, reliability of older adults was higher than that of young adults. ^*^*p* < 05. ^***^*p <* .001.

### Grégoire et al. (2022)

All three experiments in Grégoire et al. (2022) incorporated a paradigm that required participants to search for a unique shape singleton (circle among diamonds or diamond among circles), requiring participants to engage in singleton-search mode in the presence of color singleton distractors. Data from Experiments 1 and 2 were of particular interest given that reduced processing of valent (reward- and threat-related) distractors relative to neutral distractors was observed in these experiments whereas the opposite was observed in Experiment 3, although reliabilities from all three experiments are reported for completeness. Data from both the training and test phases of each experiment were combined given that mechanisms of attention capture by the distractor were identical in both phases and the only difference in the test phase was the absence of feedback, which provided sufficient data to conduct a split-half analysis. Over all experiments, the critical distractor condition comparison was attention capture by the reward (Experiments 1-3) or threat-related distractor (Experiment 1 only) vs. the neutral distractor.

When comparing the difference in oculomotor measures between the threat-related vs. neutral distractor in Experiment 1, correlation values over the measure of oculomotor capture was *r* = .609 and over fixation time was *r* = .419. Like in Kim and Anderson (2022), we found that the reliability of the learning-dependent reduction in oculomotor capture was significantly superior compared to that observed using fixation time, *p <* .001. When comparing oculomotor measures between the reward-related vs. neutral distractor, the correlations between the critical distractor conditions over oculomotor capture were *r =* .692 and *r =* .658, and over fixation time were *r =* .351 and *r =* .372, across Experiments 1 and 2, respectively. Using non-parametric randomization tests, we again found that the reliability of the learning-dependent reduction in oculomotor capture was significantly superior compared to that observed using fixation time across both Experiments, *ps <* .001 (see Figure 1). Similar results were obtained in the context of oculomotor capture in the third experiment, although overall reliability was somewhat reduced (*r* = .492 for oculomotor capture and *r* = .272 for fixation time, *p* < .001)

### Kim et al. (2024)

In Experiment 1 of Kim et al. (2024), the task required searching for a specific target shape (circle or diamond, counterbalanced across participants), requiring participants to engage in feature-search mode, which generally promotes the suppression of salient distractors (Gaspelin et al., 2015, 2017; Gaspelin & Luck, 2018). We compared trials in which a salient color singleton distractor was present vs. absent (equally often) and separately for young adults (18-23 years old) and older adults (51-79 years old). Given that we measured attention capture by first fixations to the distractor on distractor-present trials, we summed the first fixations on non-targets in distractor-absent trials and divided the total by the number of non-targets in the visual search array to calculate the probability of fixating at any one non-target (proxy distractor on distractor-absent trials). When comparing oculomotor measures between these distractor conditions, correlations over oculomotor capture (probability of fixating a [proxy] distractor on distractor present vs. absent trials) were *r* = .656 for young adults and *r =* .765 for older adults while correlations over fixation times (latency to fixate the target on distractor present vs. absent trials) was *r* = .547 for young adults and *r* = .586 for older adults. Both young and older adults demonstrated superior reliability for oculomotor capture compared to fixation times, *ps <* .001 (see Figure 1). In addition, older adults demonstrated superior oculomotor capture reliability compared to young adults, *p* < .001 (see Figure 1). However, fixation time reliability was not significantly different between age groups, *p* = .229.

In Experiment 2, the task required searching for a unique shape singleton (circle among diamonds or diamond among circles) necessitating participants to engage in singleton-search mode. Under these conditions, attentional capture by the color singleton distractor is robust and difficult to suppress, requiring reactive distractor disengagement to complete the task (Bacon & Egeth, 1994; Geng, 2014; Theeuwes, 1992; Theeuwes et al., 1998). Again, we compared trials in which the distractor was present vs. absent (equally often) and separately for young adults (19-30 years old) and older adults (57-80 years old). When comparing oculomotor measures between these distractor conditions, correlations over oculomotor capture were *r* = .815 for young adults and *r =* .890 for older adults while correlations over fixation times were *r* = .540 for young adults and *r* = .693 for older adults. As in Experiment 1, both young and older adults demonstrated superior reliability for oculomotor capture compared to fixation times, *ps <* .001 (see Figure 1). Furthermore, older adults demonstrated superior oculomotor capture reliability compared to young adults, *p* = .016, in addition to superior fixation time reliability, *p* < .001 (see Figure 1).

## Discussion

Our findings demonstrate that, as a measure, oculomotor capture produces superior reliability compared to measures computed from fixation time across numerous critical distractor comparisons. Using the probability of fixating the distractor, reliable learning-dependent reductions in distractor processing can be observed (Grégoire et al., 2022; Kim & Anderson, 2022), in addition to a measure of attention capture that is reliable for both young and older adults regardless of whether capture is overall suppressed under conditions of feature-search vs. singleton-search. Even when accounting for the increased variance in difference score calculations (Miller & Ulrich, 2013; Paap & Sawi, 2016; Weichselbaum et al., 2018), we demonstrate that oculomotor measures of attention capture on average exhibit strong reliability (mean across acquired values, *r =* .735) and are considerably more reliable than response time-based measures (Anderson & Kim, 2019; Freichel et al., 2023; Garre-Frutos et al., 2024; Ivanov et al., 2023).

Experimental psychologists have largely undervalued the utility of individual differences, and relationships between mechanisms of attentional control and other cognitive or self-report measures have been relatively unexplored. However, researchers investigating working memory capacity have examined individual differences to identify interactions between neural networks of memory and attention. Prior findings reveal that individuals with low working memory capacity exhibited stronger value-driven attentional capture (Anderson et al., 2011) and also took longer to disengage attention from a task-irrelevant distractor (Fukuda & Vogel, 2011). This relationship between working memory and attention is thought to be mediated by the locus coeruleus-noradrenaline system, particularly through modulation of the fronto-parietal attention networks (Unsworth & Robison, 2017). However, individual differences in working memory capacity were unable to predict performance in visual search tasks requiring feature or conjunction search (Kane et al., 2006). The lack of a relationship here is informed by the findings of Ivanov et al. (2023) in which attention capture and learning-dependent distractor suppression were investigated as potentially useful measures of individual differences using manual response times. Unfortunately, both within- and between-session reliability for both measures were poor despite robust group level differences across conditions, suggesting that inconsistent findings relating individual differences in working memory capacity to attention may be due in part to the use of measures with poor reliability (all of the aforementioned studies and many similar studies used attention measures derived from manual response times). Interestingly, when value-driven attentional capture was measured from distractor fixations (Anderson & Yantis, 2012), the reported correlation with working memory capacity was numerically quite a bit stronger than when value-driven attentional capture was measured from manual response times (Anderson et al., 2011). Our findings suggest a potential path toward more consistent outcomes relating attention measures to other cognitive processes like working memory, and to the more fruitful exploration of individual differences in the learning-dependent control of attention more generally through fixation-based measures of attentional selection. More reliable measures of attentional control are of particular importance if the goal is to predict the progression of neurodegenerative diseases and other clinical outcomes, and our findings point to the value of eye tracking in the pursuit of such measures.

The set of experiments in Kim et al. (2024) additionally revealed that older adults exhibit greater reliability compared with young adults. Older adults generally have slower response times compared with young adults and this becomes problematic as overall slower response times have greater variability (Kim et al., 2024; Tse et al., 2010). Although Experiment 2 demonstrated that older adults make more first fixations to the distractor compared with young adults, superior reliability cannot be reduced to a product of this greater capture effect given that Experiment 1 showed similar oculomotor suppression by the distractor in both age groups but still greater reliability in older adults. The strong reliability of oculomotor measures in older adults can address a significant issue in the aging literature of low reliability due to increased error variance in measures like response time. Furthermore, the relatively higher reliability in Kim et al. (2024) suggests that the reliability of salience-driven capture may be higher compared with statistically learned distractor suppression (Grégoire et al., 2022; Kim & Anderson, 2022), which is in line with the results of Ivanov et al. (2023).

A natural question posed by the findings of the present study is why oculomotor capture produces a more reliable measure of distractor processing than fixation time in addition to what is typically observed in the literature with respect to manual response time. Although we can only speculate, this superior reliability may be found in the ballistic nature of the measure.

Oculomotor capture essentially measures the probability that a task-irrelevant stimulus evokes greater attentional priority than the target at the time of saccade initiation, being directly linked to distractor-target competition in the visual system. Manual response time-based measures add a host of post-selection processes that are tied to target-response mappings and the execution of a manual response (often a keypress), all of which contribute variability that is removed when assessing oculomotor capture. Even in the context of fixation time, the time required to disengage attention from any non-target that is fixated and the efficiency with which the subsequent eye movement is targeted contribute additional variability that occurs after oculomotor capture is assessed, during which there is additional opportunity for task-unrelated processes (e.g., mind wandering) to randomly slow responses. If the goal is to measure distractor processing, the probability of initially fixating the distractor (oculomotor capture) may be the purest and most direct means of assessing it.

Our findings across multiple experiments suggests that the superior reliability of oculomotor capture relative to even response time-based measures derived from eye tracking may reflect a more general property of the measurements that would further generalize to other tasks and experimental situations. However, determining whether this is the case requires further investigation, in addition to the extent to which specific mechanisms of distractor processing (e.g., learning effects that promote capture vs. suppression, salience-driven vs. learning-dependent priority) are differently reliable. Similarly, it would also be important to investigate whether the observed high reliability of oculomotor capture as a measure extends to other mechanisms of distractor processing (e.g., contingent attention capture, emotion-modulated distraction).

The present study suggests a potential avenue forward for the field of psychological science to maximize reproducibility by utilizing oculomotor measures that exhibit high reliability. However, the biggest limitation in acquiring such measures is the accessibility of eye tracking technology. All of the datasets analyzed utilized an EyeLink 1000 plus eye tracker (SR Research) that is far less accessible than what is required to conduct research using manual response time measures, both with respect to financial cost and training. The development of more reliable measures of visual information processing involving manual response time that can more closely approximate what we were able to achieve with oculomotor measures is therefore an important target for future research. At least for the time being, until more reliable response time-based measures are developed, we recommend that researchers consider investing in oculomotor measures particularly when individual differences in distractor processing are of scientific interest. Oculomotor measures are naturally bound to experiments involving the processing of visual information, and it is also important to identify reliable measures of information processing in other sensory modalities in an effort to maximize statistical power and reproducibility.

